# A Fluorescent Dauer Marker in *Caenorhabditis inopinata* Enables Comparative Analysis of Dauer-Inducing Mechanisms

**DOI:** 10.64898/2026.04.06.716796

**Authors:** Ryo Iitsuka, Shun Oomura, Nami Haruta, Asako Sugimoto

**Affiliations:** Laboratory of Developmental Dynamics, Graduate School of Life Sciences, Tohoku University, Sendai 980-8577, Japan

**Keywords:** dauer, *Caenorhabditis elegans*, *Caenorhabditis inopinata*, fluorescent marker

## Abstract

Dauer larvae are a dormant developmental stage in nematodes that is induced by a range of environmental cues. The molecular mechanisms that transduce these cues to regulate dauer entry have been well characterized in *Caenorhabditis elegans*, whereas those in other nematode species remain unclear. The closest known sibling species of *C. elegans, Caenorhabditis inopinata*, occupies a distinct ecological niche and shows an extremely low frequency of dauer formation by starvation in laboratory conditions, suggesting that it could serve as a useful comparative model for analyzing dauer-inducing mechanisms. To support such analysis, we generated a fluorescent dauer reporter, *Cin-col-183p::mCherry*, in *C. inopinata* based on a previously reported dauer-specific reporter in *C. elegans*. This reporter showed fluorescence specifically in the pre-dauer and dauer stages, but not in other developmental stages, indicating that it functions as a dauer-specific marker in *C. inopinata*. Using these marker strains, we compared the responses to high temperature and RNAi-mediated knockdown of insulin/IGF-1 pathway genes (*daf-2, age-1*, and *pdk-1*), and found that dauer induction differs mechanistically between *C. elegans* and *C. inopinata*. This dauer-specific fluorescent strain will be a useful tool for investigating the diversity of dauer-inducing mechanisms across nematode species.

**Article Summary:** Dauer is a dormant developmental stage in nematodes induced by environmental stress. Although its regulation is well studied in *Caenorhabditis elegans*, the mechanisms in other species remain unclear. Here, we developed a fluorescent dauer reporter, *Cin-col-183p::mCherry*, in *Caenorhabditis inopinata*, a close relative of *C. elegans*. The reporter was specifically expressed in pre-dauer and dauer stages, confirming its usefulness as a dauer marker. Using this strain, we found that responses to high temperature and insulin/IGF-1 pathway gene knockdown differ between *C. elegans* and *C. inopinata*. This reporter will help reveal diversity in dauer-inducing mechanisms across nematode species.

## Introduction

Dormancy is a physiological state characterized by reduced metabolic activity that allows organisms to survive adverse environmental conditions. In the phylum Nematoda, the dormant stage is known as the dauer larva, an adaptive developmental stage that enables survival under environmental stress, typically without feeding (McSorley, 2003; Vlaar et al., 2021). Entry into the dauer stage occurs through various mechanisms among nematodes. These include environmental cues, such as starvation, high temperature, and vector-associated signals, as well as developmentally programmed processes (Ailion and Thomas, 2000; Cassada and Russell, 1975; Roeber et al., 2013; Zhao et al., 2013). These differences in dauer entry raise the question of the extent to which the underlying mechanisms of dauer formation are conserved or diversified across the phylum Nematoda.

The mechanisms of dauer entry in nematodes have been extensively studied in *Caenorhabditis elegans*. In *C. elegans*, dauer formation is an alternative third larval stage (L3) triggered by harsh environmental conditions such as starvation, high temperatures, and pheromone (Ailion and Thomas, 2000; Cassada and Russell, 1975; Golden and Riddle, 1982) (Fig. 1a). Environmental cues initially induce formation of pre-dauer larvae, an alternative L2 stage, and then the larvae can commit to dauer stage under continued unfavorable conditions (Golden and Riddle, 1984) (Fig. 1a). Dauer larvae are morphologically distinct from reproductive stages, characterized by a thin body, a thickened cuticle with lateral alae, and resistance to 1% SDS (Cassada and Russell, 1975).

**Fig. 1.**
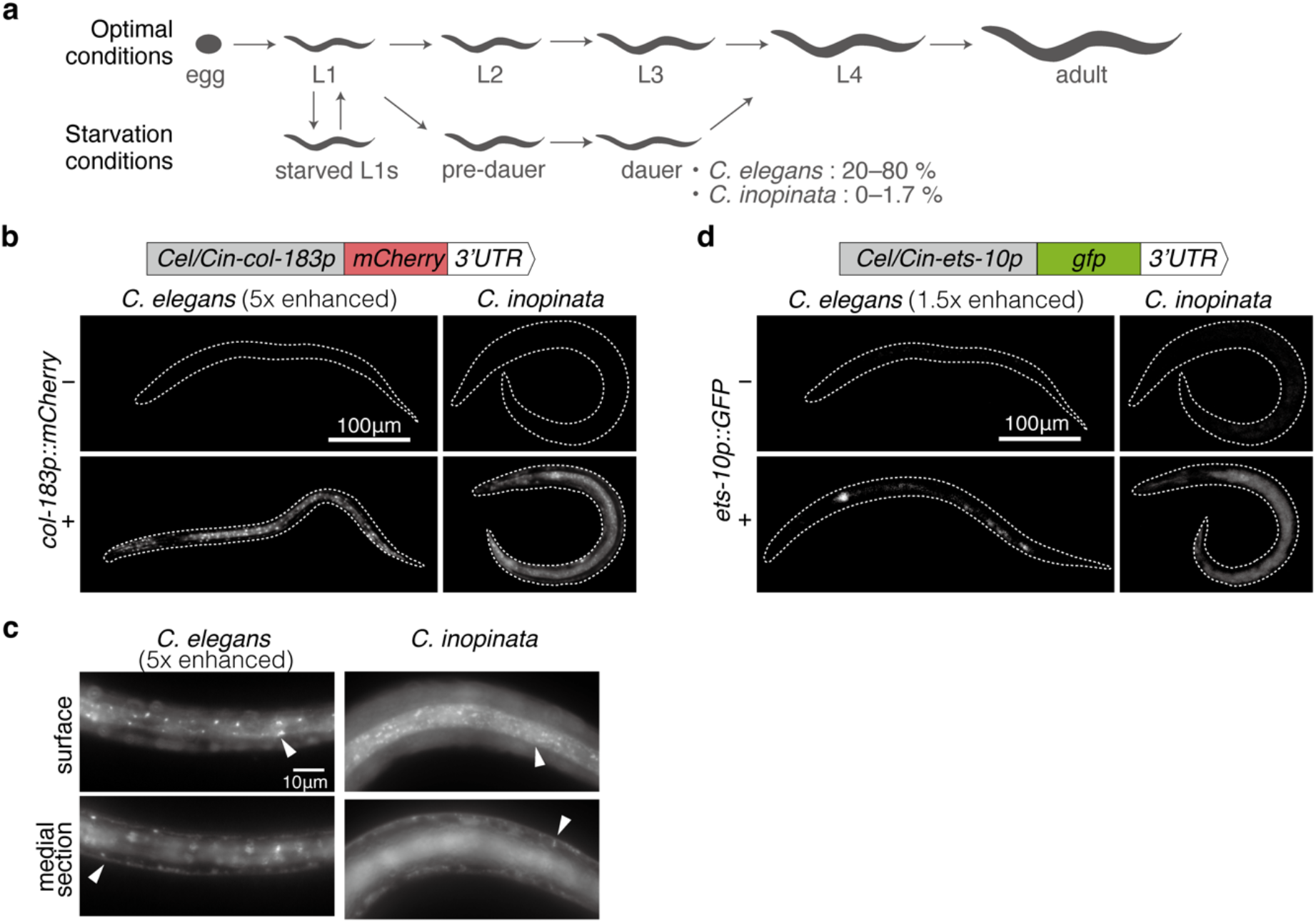
Establishment of transgenic lines harboring dauer-specific reporters in *C. elegans* and *C. inopinata*. a) Schematic diagram illustrating the developmental trajectories of *C. elegans* and *C. inopinata* under optimal and starvation conditions. Under optimal conditions, animals develop from the egg through larval stages (L1–L4) to adulthood. Under starvation conditions, animals either arrest at the L1 stage (starved L1s) or enter the pre-dauer stage, followed by the dauer stage. Reported dauer frequency in both species under laboratory starvation conditions is indicated (Cassada and Russell, 1975; Hammerschmith et al., 2022). b) *Cel*/*Cin-col-183p::mCherry* transgenic strains. The transgene constructs and fluorescence images of whole non-transgenic (top) and transgenic (bottom) dauer larvae are shown. c) Magnified fluorescence micrographs of the surface and medial sections of the lateral midbody regions of *Cel*/*Cin-col-183p::mCherry* dauer larvae. Arrowheads indicate fluorescence in the hypodermal-like tissues. d) *Cel/Cin-ets-10p::GFP* transgenic strains. The transgene constructs and fluorescence images of whole non-transgenic (top) and transgenic (bottom) dauer larvae are shown. Scale bars: 100 μm (b, d) and 10 μm (c). Fluorescence intensity in *C. elegans* was enhanced 5-fold in (b,c) and 1.5-fold in (d), relative to that in *C. inopinata* for visualization. Dotted outlines indicate worm bodies in (b, d).

Genetic analyses in *C. elegans* have shown that four conserved signaling pathways—the cGMP, insulin/IGF-1, TGF-β, and steroid hormone biosynthesis pathways—are involved in dauer formation. These pathways are proposed to function as a regulatory cascade, in which the cGMP pathway primarily senses environmental cues and transmits signals to the insulin/IGF-1 and TGF-β pathways. These, in turn, regulate the steroid hormone biosynthesis pathway, which ultimately determines whether dauer formation or reproductive development occurs (Fielenbach and Antebi, 2008). Although genes involved in these four signaling pathways are broadly conserved across nematodes (Gilabert et al., 2016; Vlaar et al., 2021), their specific roles in dauer formation in other nematode species remain unclear.

*Caenorhabditis inopinata* is the species most closely related to *C. elegans* (Kanzaki et al., 2018); however, their life cycles, including the induction of dauer formation, differ markedly. Whereas *C. elegans* inhabits various types of rotting plant material (Félix and Braendle, 2010), *C. inopinata* occupies a specialized niche within the syconia of the fig *Ficus septica*. The optimal culture temperature of *C. inopinata* (25–29°C) is higher than that of *C. elegans* (20°C). Under laboratory starvation conditions, the frequency of dauer formation in *C. inopinata* is extremely low (0–1.7 %) (Hammerschmith et al., 2022) compared with that in *C. elegans* (20–80 %) (Cassada and Russell, 1975). These differences in habitat and in responses to starvation suggest that the mechanisms underlying dauer induction may differ between the two species. Therefore, *C. inopinata* and *C. elegans* provide an excellent system for comparative analyses aimed at understanding the conservation and diversification of dauer formation mechanisms.

In this study, to facilitate a comparative analysis of dauer-inducing mechanisms between *C. inopinata* and *C. elegans*, we developed a fluorescent dauer reporter, *Cin-col-183p::mCherry*, in *C. inopinata* based on a previously established dauer-specific reporter in *C. elegans* (Shih et al., 2019). This *C. inopinata* reporter strain enables sensitive detection of dauer entry and efficient comparative assessment of dauer phenotypes. Furthermore, using the *Cin-col-183p::mCherry* marker, we demonstrated that the contributions of temperature and insulin/IGF-1 pathway genes (*daf-2, age-1*, and *pdk-1*) to dauer induction in *C. inopinata* differ from those in *C. elegans*.

## Results and Discussion

### Identification of *C. inopinata* orthologs of the dauer-specific genes in *C. elegans*

To efficiently detect the dauer state in *C. inopinata*, we aimed to construct dauer-specific fluorescent reporter strains based on the previously reported dauer-specific fluorescent reporters in *C. elegans* (Shih et al. 2019). In that study, promoters of the *C. elegans* dauer-specific genes (*col-183, ets-10*, and *nhr-246*) were used to construct fluorescent reporters. In *C. elegans, col-183* encoding a collagen is exclusively expressed in the hypodermis during the pre-dauer and dauer stages; *ets-10* encoding a ETS-class transcription factor is expressed in the intestine, some neurons, and weakly in the hypodermis during the pre-dauer and dauer stages, as well as in the uterine and spermathecal cells during the L4 and adult stages; *nhr-246* encoding an nuclear hormone receptor is expressed strongly in the intestine and hypodermis during pre-dauer and dauer stages (Shih et al. 2019). Orthologs of these genes in *C. inopinata* were identified by reciprocal BLASTP searches (hereafter, the prefixes “*Cel-”* or “*Cin-*” are used to distinguish *C. elegans* and *C. inopinata* orthologs, respectively). For *Cel-col-183* and *Cel-ets-10*, one-to-one orthologs (*Cin-col-183* and *Cin-ets-10*, respectively) were identified, and their promoter regions were highly conserved relative to those of their *C. elegans* counterparts. In contrast, the top hit for *Cel-nhr-246* in *C. inopinata* corresponded to multiple homologs in *C. elegans*. We therefore focused on *Cin-col-183* and *Cin-ets-10* as candidate dauer-specific genes in *C. inopinata*.

### Construction of transgenic lines harboring dauer-specific marker transgenes in *C. inopinata*

We constructed transgenes expressing the fluorescent proteins mCherry and GFP under the control of the *Cel/Cin-col-183* and *Cel/Cin-ets-10* promoters, respectively (Fig. 1b, d). The *C. elegans* transgenes were integrated into the *C. elegans* genome using the miniMos method (Frøkjær-Jensen et al., 2014), whereas the *C. inopinata* transgenes were integrated into the *C. inopinata* genome by microparticle bombardment (Oomura et al., 2022).

We examined fluorescent protein expression from the transgenes in starvation-induced dauer larvae (Fig. 1b–d). In *C. inopinata* dauer larvae, *Cin-col-183p::mCherry* showed strong fluorescence in hypodermis along the longitudinal ridges, similar to the pattern previously reported in *C. elegans* dauer larvae (Shih et al., 2019) (Fig. 1b, c). The fluorescence of both *Cel-col-183p::mCherry* and *Cin-col-183p::mCherry* was sufficiently bright to be detected under a fluorescent stereomicroscope (Fig. 2b). Although *Cin-ets-10p::GFP* showed fluorescence in the intestine of dauer larvae, resembling the previously reported pattern in *C. elegans* (Shih et al., 2019), its signal intensity was insufficient for reliable detection under a stereomicroscope (Fig. 1d). We therefore focused subsequent analysis on *Cin-col-183p::mCherry*.

**Fig. 2.**
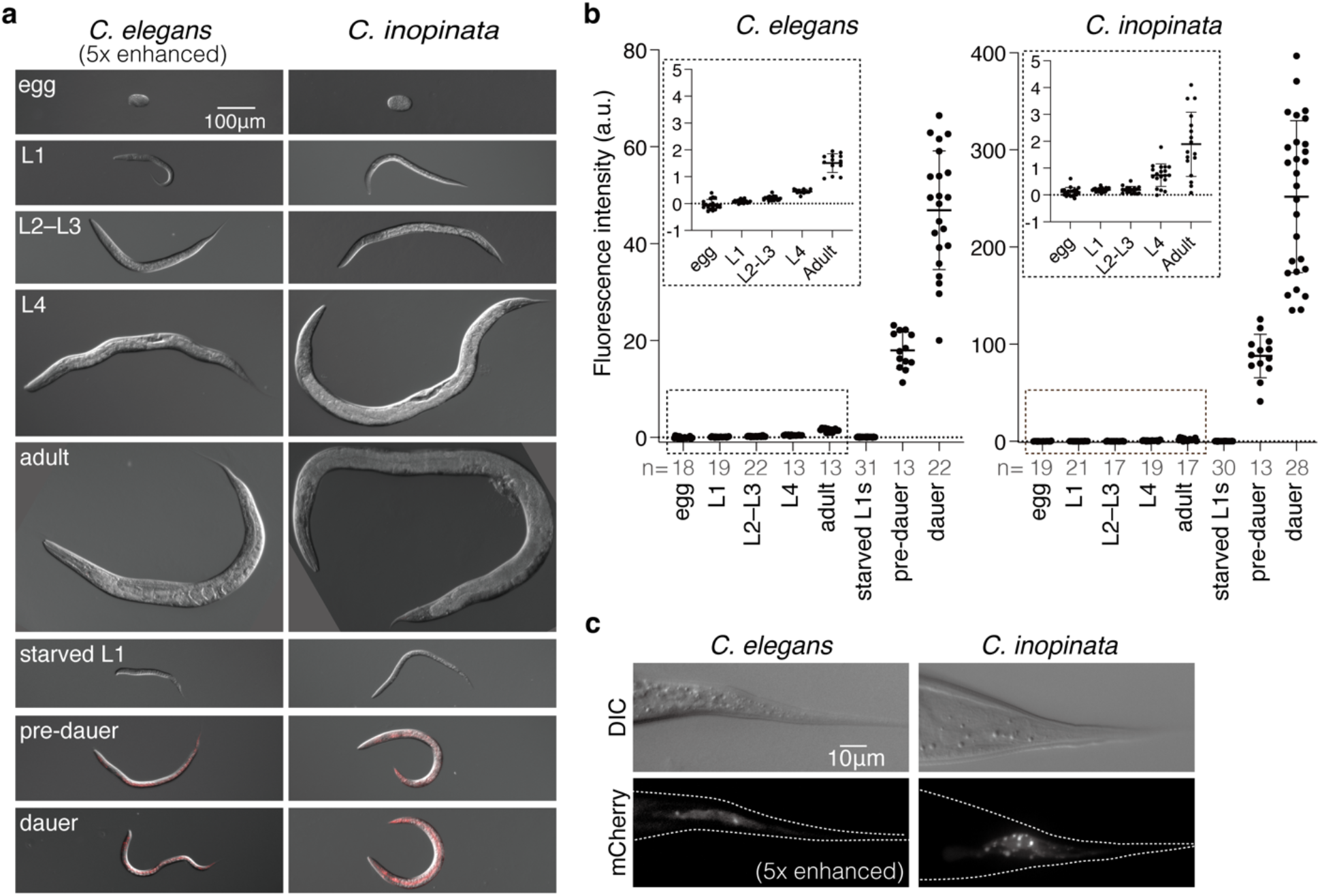
Developmental stage-specific expression of the *Cel/Cin-col-183p::mCherry* markers. a) Representative merged DIC and mCherry fluorescence (red) images of *Cel-col-183p::mCherry* (left) and *Cin-col-183p::mCherry* (right) under optimal conditions (egg, L1, L2–L3, L4, and adult), and starvation conditions (starved L1, pre-dauer, and dauer). Scale bar: 100 μm. b) Quantification of fluorescence intensity at each stage. Insets show the boxed regions at a larger scale. Each dot represents an individual animal. Bars indicate mean ± SD. n = the number of animals analyzed. c) Magnified views of the tail regions of an adult *C. elegans* hermaphrodite (left) and an adult *C. inopinata* female (right). Dotted outlines indicate worm bodies. Scale bar: 10 μm. In a) and c), mCherry fluorescence intensity in *C. elegans* was enhanced 5-fold relative to that in *C. inopinata* for visualization.

### *Cin-col-183p::mCherry* is specifically expressed after dauer entry in *C. inopinata*

*Cin-col-183p::mCherry* fluorescence was examined across developmental stages. In *C. inopinata*, fluorescence remained at background levels throughout the reproductive stages from egg to adult, except for the tail tip fluorescence in the L4 and adult stages, similar to that reported previously in *C. elegans* (Shih et al., 2019) (Fig. 2, mean fluorescence intensity during reproductive stages ranged from −0.29 to 1.95 a.u. (arbitrary unit) in *C. elegans* and from −0.13 to 4.29 a.u. in *C. inopinata*). The tail tip fluorescence (Fig. 2c) is likely derived from hypodermal cells (Nguyen et al., 1999), allowing it to be distinguished from the broad hypodermal expression observed in dauer larvae.

To determine whether *Cin-col-183p::mCherry* is expressed specifically after dauer entry or in response to starvation alone, we next quantified fluorescence under starvation conditions, namely in pre-dauer larvae, dauer larvae, and starvation-induced L1-arrested larvae (starved L1s). The pre-dauer stage is a starvation-induced alternative L2 stage that precedes dauer formation; larvae at this stage resemble dauer larvae morphologically but remain sensitive to 1% SDS, unlike dauer larvae (Fig. 1a) (Golden and Riddle, 1984). Starved L1s are developmentally arrested L1 larvae that have experienced starvation (Fig. 1a) (Johnson et al., 1984).

In dauer larvae, *Cel-col-183p::mCherry* and *Cin-col-183p::mCherry* were strongly expressed in hypodermis in both *C. elegans* and *C. inopinata*, respectively, and the mean fluorescence intensity was approximately five-fold higher in *C. inopinata* than in *C. elegans* (Fig. 2a, b, mean of fluorescence intensity: 46.9 a.u. in *C. elegans* and 251.8 a.u. in *C. inopinata*). Pre-dauer larvae of both species exhibited weaker fluorescence than dauer larvae but stronger than animals in reproductive stages (mean of fluorescence intensity: 18.0 a.u. in *C. elegans* and 87.9 a.u. in *C. inopinata*). In contrast, fluorescence intensities in starved L1s were as low as those in reproductive stages in both species (mean of fluorescence intensity: 0.07 a.u. in *C. elegans* and 0.27 a.u. in *C. inopinata*). These results indicate that *Cin-col-183p::mCherry* expression is not induced by starvation alone, but is associated with dauer entry (Fig. 2a, b).

Together, these results demonstrate that, like *Cel-col-183p::mCherry* in *C. elegans* (Shih et al., 2019), *Cin-col-183p::mCherry* is specifically expressed in both pre-dauer and dauer stages and can therefore serve as a reliable and sensitive marker of dauer entry in *C. inopinata*.

### Regulatory mechanisms of dauer induction differ between *C. elegans* and *C. inopinata*

Using *Cin-col-183p::mCherry* as a dauer entry marker, we asked whether high temperature can trigger dauer entry in *C. inopinata*. In *C. elegans*, high temperature (27°C) induces dauer formation at a low frequency (0–20%) (Ailion and Thomas, 2000). Consistent with this, in the *Cel-col-183p::mCherry* strain that we constructed, fluorescence was undetectable at 20°C, whereas at 27°C, a subset of L4 and adult animals showed strong *Cel-col-183p::mCherry* fluorescence, implying partial activation of the dauer program (Fig. 3). Because *C. inopinata* grows at a higher temperature range (25°C–31°C; Supplementary Fig. 2) than *C. elegans* and has an optimal temperature is 27°C (Kanzaki et al., 2018; Woodruff et al., 2019), 31°C was used as the high-temperature condition. In contrast to *C. elegans, C. inopinata* showed no detectable *Cin-col-183p::mCherry* fluorescence and developed to L4 or adult stages at both 27°C and 31°C (Fig. 3), suggesting that the temperature responses of dauer entry differ between the two species.

**Fig. 3.**
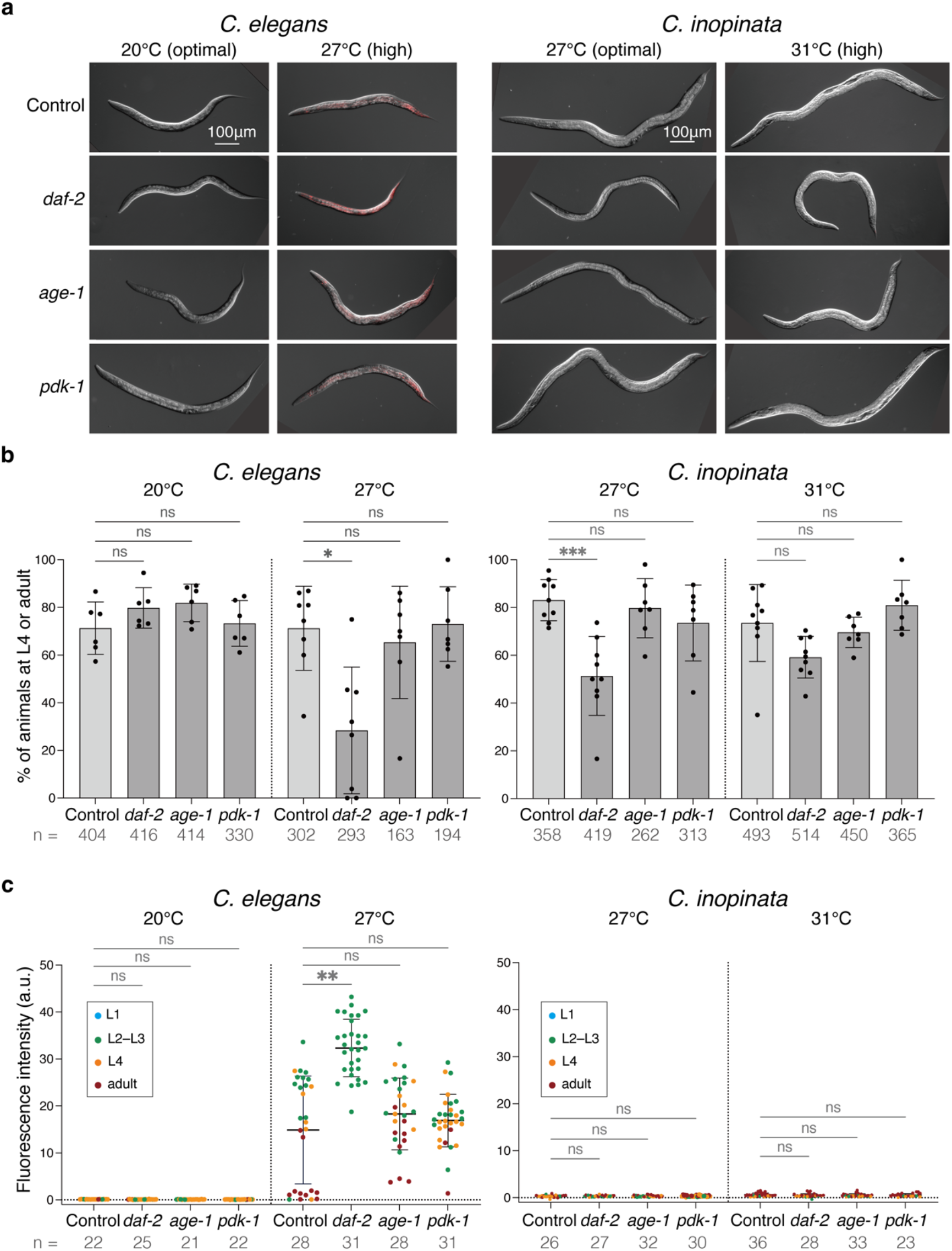
Distinct responses to dauer-inducing signals measured by *Cel/Cin-col-183p::mCherry* markers. a) Representative merged DIC and mCherry fluorescence (red) images of control and RNAi-treated animals under optimal and high temperature conditions. Images were acquired after incubation for 48 h at 20°C or 72 h at 27°C for the *Cel-col-183p::mCherry* strain, and for 72 h at 27°C or 31°C for the *Cin-col-183p::mCherry* strain. Scale bar: 100 μm. b) Proportion of animals that reached the L4 or adult stages under each condition. Each data point represents a single plate. c) Quantification of the mCherry fluorescence intensity. Each dot represents an individual animal. Dot colors indicate developmental stages: L1 (blue), L2–L3 (green), L4 (orange), and adult (dark red). In b) and c), bars indicate mean ± SD. n = the number of animals analyzed. Statistical significance was evaluated within each species and each temperature condition using the Kruskal-Wallis test followed by Dunn’s multiple comparisons test against the control group; \**P* < 0.05, \*\**P* < 0.01, \*\*\**P* < 0.001, ns: not significant.

We next compared the effects of the insulin/IGF-1 pathway genes (*Cel*-*daf-2/InsR, Cel*-*age-1/PI3K*, and *Cel-pdk-1/PDK*) to dauer entry at high temperature. In *C. elegans*, reduced function of these genes enhances dauer entry (Dillin et al., 2002; Paradis et al., 1999). Consistent with previous reports, RNAi of *Cel-daf-2* at 27°C caused L3-like developmental arrest with the *Cel-col-183p::mCherry* fluorescence at a higher frequency than the control (Fig. 3, Supplementary Fig. 3). RNAi of *Cel-age-1* or *Cel-pdk-1* at 27°C also increased the proportion of *Cel-col-183p::mCherry* positive animals, although most animals developed to the L4s and adult stage, indicating partial activation of the dauer program (Fig. 3). In *C. inopinata*, however, all animals treated with *Cin-daf-2, Cin-age-1*, and *Cin-pdk-1* RNAi at 31°C developed to adulthood without detectable *Cin-col-183p::mCherry* fluorescence (Fig. 3). Instead, *Cin-daf-2* RNAi caused a significant developmental delay at 27°C (Fig. 3b, Supplementary Fig. 3). Together, these results indicate that, unlike *C. elegans*, reduced function of *Cin-daf-2, Cin-age-1*, and *Cin-pdk-1* does not activate the dauer program indicated by the *col-183* reporter expression in *C. inopinata* even at high temperature.

Thus, in addition to the response to starvation (Hammerschmith et al., 2022), the responses of *C. inopinata* to high temperature and to perturbation of insulin/IGF-1 pathway genes also differ from those of *C. elegans*. These findings indicate that the environmental and genetic regulation of dauer entry in *C. inopinata* is distinct from that in *C. elegans*.

## Conclusion

In this study, we established a fluorescent reporter, *Cin-col-183p::mCherry*, that enables the sensitive detection of pre-dauer and dauer larvae in *C. inopinata*. This reporter is expected to facilitate investigation of the dauer-inducing mechanisms in *C. inopinata*, which appear to differ from those in *C. elegans*, as indicated by the distinct responses to starvation, high temperature, and inhibition of the insulin/IGF-1 signaling pathway. For example, the *Cin-col-183p::mCherry* strain enables efficient screening for genes involved in dauer signaling pathways, as well as exploration of environmental cues that induce dauer formation in *C. inopinata*. These applications will contribute to a broader understanding of the diverse dauer-inducing mechanisms across nematodes.

## Materials and methods

### *C. elegans* and *C. inopinata* maintenance

*C. elegans* strains were maintained on Nematode Growth Medium (NGM) plates seeded with *Escherichia coli* strain OP50 as a food source (Brenner, 1974). *C. inopinata* strains were maintained on NGM plates with a higher agar concentration to prevent burrowing, and were fed *Escherichia coli* HT115 (DE3) (Oomura et al., 2022). The strains used and constructed in this study are listed in Supplementary Table 1.

### Identification of *C. inopinata* orthologs of *C. elegans* dauer-specific genes

*C. inopinata* orthologs of *C. elegans* dauer-specific genes, *Cel-col-183, Cel-ets-10*, and *Cel-nhr-246*, were identified by reciprocal BLASTP searches. Each *C. elegans* protein was used as a query against the *C. inopinata* protein database (PRJDB5687) using BLASTP on WormBase Parasite (https://parasite.wormbase.org/), and the top-hit proteins were then searched back against the *C. elegans* protein database (PRJNA13758) using BLASTP on WormBase (https://www.wormbase.org/). *Cin-col-183* (WormBase Parasite gene ID: Sp34_X0149100) and *Cin-ets-10* (gene ID: Sp34_X0258400) were identified as one-to-one orthologs, whereas the top hit for *Cel-nhr-246* in *C. inopinata* corresponded to multiple homologs in *C. elegans*.

The promoter regions of *Cin-col-183* and *Cin-ets-10* were identified by aligning their 5’ upstream regions with the corresponding promoter regions of *Cel-col-183* (1695 bp) and *Cel-ets-10* (1111 bp) used in Shih et al. (2019), using Pro-Coffee (https://tcoffee.crg.eu/apps/tcoffee/do:procoffee). The highly conserved regions, *Cin-col-183p* (1877 bp) and *Cin-ets-10p* (1047 bp), were used as promoters for transgene construction.

For RNAi analysis, we used the *C. inopinata* one-to-one orthologs of *Cel-daf-2, Cel-age-1*, and *Cel-pdk-1* proposed under “Orthologues” on WormBase Parasite (gene ID: Sp34_30289300, Sp34_ 20327500, Sp34_X0009000, respectively).

### Plasmid construction and transgenesis

All plasmids used to construct dauer-specific reporters were generated by three-fragment Gibson assembly (Gibson et al., 2009) into pTK68 (Oomura et al., 2022), which contains a hygromycin B-resistant cassette (*Cel-rps-0p::hygB::Cel-unc-54 3’UTR*). The plasmid pRI1, containing the *Cel-col-183p::mCherry* transgene, and pRI3, containing the *Cin-col-183p::mCherry* transgene, were constructed by assembling three fragments into SacII/ApaI digested pTK68: the *Cel/Cin-col-183* promoter regions, the *mCherry* ORF amplified from pMTN1R (Toya et al., 2010), and the *Cel-unc-54 3’UTR* amplified from pTK68. Similarly, plasmids pRI2 and pRI4, containing *Cel -ets-10p::GFP* and *Cin-ets-10p::GFP*, respectively, were constructed using the *Cel*/*Cin-ets-10* promoter regions, the GFP ORF amplified from pMTN1G (Toya et al., 2010), and the *Cel-unc-54 3’UTR*. Promoter fragments were amplified from the genomic DNA of each species. All primers used in this study are listed in Supplementary Table 2.

Plasmids pRI1 or pRI2 were integrated into the wild-type *C. elegans* strain N2 using the miniMos method (Frøkjær-Jensen et al., 2014). The single copy insertions of *Cel-col-183p::mCherry* and *Cel-ets-10p::GFP* were mapped on Chr. I: 4,995,933 and Chr. V: 3,035,195, respectively, by inverse PCR.

Transgenic *C. inopinata* lines were generated using microparticle bombardment (Oomura et al., 2022). Plasmids pRI3 and pRI4 were co-bombarded into the wild-type *C. inopinata* strain NKZ35, and transformants were selected based on hygromycin B resistance (Oomura et al., 2022). Two independent lines were established: SA1660, which was positive for both *Cin-col-183p::mCherry* and *Cin-ets-10p::GFP* fluorescence, and SA1664, which was positive only for *Cin-col-183p::mCherry* fluorescence.

### Staging of worms

Worm staging in this study was defined as follows. Reproductive stages (L1–L4 and adult) are worms grown under well-fed conditions. Dauer larvae are larvae induced by starvation that are resistant to 1% SDS and exhibit alae. Pre-dauer larvae are larvae induced by starvation that are similar in body length to dauer larvae but sensitive to 1% SDS (Golden and Riddle, 1984). Starved L1s are L1 larvae maintained under starvation conditions (Johnson et al., 1984).

Dauer larvae were induced by 4–6 days of bacterial depletion in *C. elegans*, whereas in *C. inopinata*, they were induced by 12–14 days of bacterial depletion. Dauer larvae were isolated as survivors of 1% SDS treatment for 15 minutes in *C. elegans* (Cassada and Russell, 1975) and 30 minutes in *C. inopinata* (Hammerschmith et al., 2022). The presence of alae in SDS-resistant larvae was confirmed by differential interference contrast (DIC) microscopy.

Pre-dauer larvae were induced by ∼3 days of bacterial depletion in *C. elegans* and ∼9 days of bacterial depletion in *C. inopinata*. The pre-dauer stage was validated by lethality following 1% SDS treatment, and siblings of the treated worms were used for subsequent analysis. Body length was measured from DIC images using ImageJ/Fiji (Mörck and Pilon, 2006) and compared with that of dauer larvae and L2–L3 larvae (Supplementary Fig. 1a).

Starved L1s were obtained as follows. Gravid adults were allowed to lay eggs on plates without food, and all adults were removed after ∼24 h. Starved L1s were collected after ∼36 h for *C. elegans* and ∼48 h for *C. inopinata* (Supplementary Fig. 1b).

Reproductive stages in *C. elegans* and *C. inopinata* were initially estimated based on the developmental timing (Byerly et al., 1976; Kanzaki et al., 2018; Woodruff et al., 2019) and then confirmed by stage-specific morphology under a stereomicroscope or a DIC microscope.

### Fluorescence microscopy and quantification analysis

DIC and fluorescence images were acquired using an Axioplan 2 imaging microscope (ZEISS) equipped with a Plan-NEOFLUAR 10x/0.30 objective lens (ZEISS) for whole-body imaging, or a Plan-APOCHROMAT 40x/0.95 objective lens (ZEISS) for imaging magnified views of dauer larvae and tail regions in adults. Animals were immobilized on 2% agarose pads with 1 mM levamisole. Fluorescence was quantified using ImageJ/Fiji software. Fluorescence intensity was measured by subtracting the background fluorescence surrounding each animal from the whole-body fluorescence.

### Feeding RNAi

All feeding RNAi experiments were performed according to the method of Timmons and Fire (1998), with the addition of 1 mM isopropyl-β-D-thiogalactopyranoside (IPTG) to the RNAi plates (Kamath et al., 2000).

To construct feeding RNAi plasmids, DNA fragments of *C. elegans* genes (*Cel-daf-2* (1531bp), *Cel-age-1* (1251bp), and *Cel-pdk-1* (435bp)) were amplified from a *C. elegans* cDNA two-hybrid library, whereas those of *C. inopinata* genes (*Cin-daf-2* (1550bp), *Cin-age-1* (1218bp), and *Cin-pdk-1* (507bp)) were amplified from *C. inopinata* genomic DNA using the primers listed in Supplementary Table 2. Each fragment was inserted into ApaI/NotI-digested T444T vector (Sturm et al., 2018) by Gibson assembly. For feeding RNAi experiments, each cloned plasmid was transformed into HT115 (DE3) competent cells. The empty vector T444T was used as a negative control, and *Cel*/*Cin-unc-22* RNAi plasmids were used as positive controls. In both *C. elegans* and *C. inopinata, Cel*/*Cin-unc-22* feeding RNAi causes a strong twitching phenotype (Kanzaki et al., 2018; Timmons and Fire, 1998).

For feeding RNAi assays, SA1692 (*Cel-col-183p::mCherry*) and SA1664 (*Cin-col-183p::mCherry*) animals were grown on HT115 (DE3) bacteria expressing each dsRNA. On each 60 mm RNAi plate, 5 gravid adults of *C. elegans* or 20 gravid adults of *C. inopinata* were allowed to lay eggs for 2–4 h before removal. Animals were scored 48 h after adults plating at 20°C for *C. elegans*, and 72 h after plating at 27°C for *C. elegans* and *C. inopinata*, or at 31°C for *C. inopinata*, corresponding to the time points at which most control animals had passed the dauer decision and reached the L4 stage (Supplementary Fig. 3).

## Data availability statement

Plasmids and strains are available upon request. The authors affirm that all data necessary for confirming the conclusions of the article are present within the article, figures, and Supplementary material.

Supplemental material available at GENETICS online.

## Acknowledgements

We thank Prof. Taisei Kikuchi and Asst. Prof. Ryusei Tanaka for providing information on the *C. inopinata* genome and for insightful discussions; Ms. Hiroko Sugawara and Ms. Makiko Sasaki for assistance with plasmid construction and nematode strain maintenance; and Dr. Ryuhei Hatanaka, Dr. Momoe Nakajo, and members of the Sugimoto laboratory for the helpful discussion. We also thank WormBase and WormBase Parasite.

## Funding

This work was supported by Japan Science and Technology Agency (JST), Core Research for Evolutionary Science and Technology (CREST) Grant Number JPMJCR18S7, Japan Society for the Promotion of Science (JSPS) KAKENHI Grant Number 23K27152 to A.S., and JST, Support for Pioneering Research Initiated by the Next Generation (SPRING), Grant Number JPMJSP2114, Advanced Graduate Program for Future Medicine and Health Care, Tohoku University to R.I.

## Conflicts of interest

The author(s) declare no conflict of interest.

## Supplementary Figures and Tables

**Supplementary Fig. 1.**
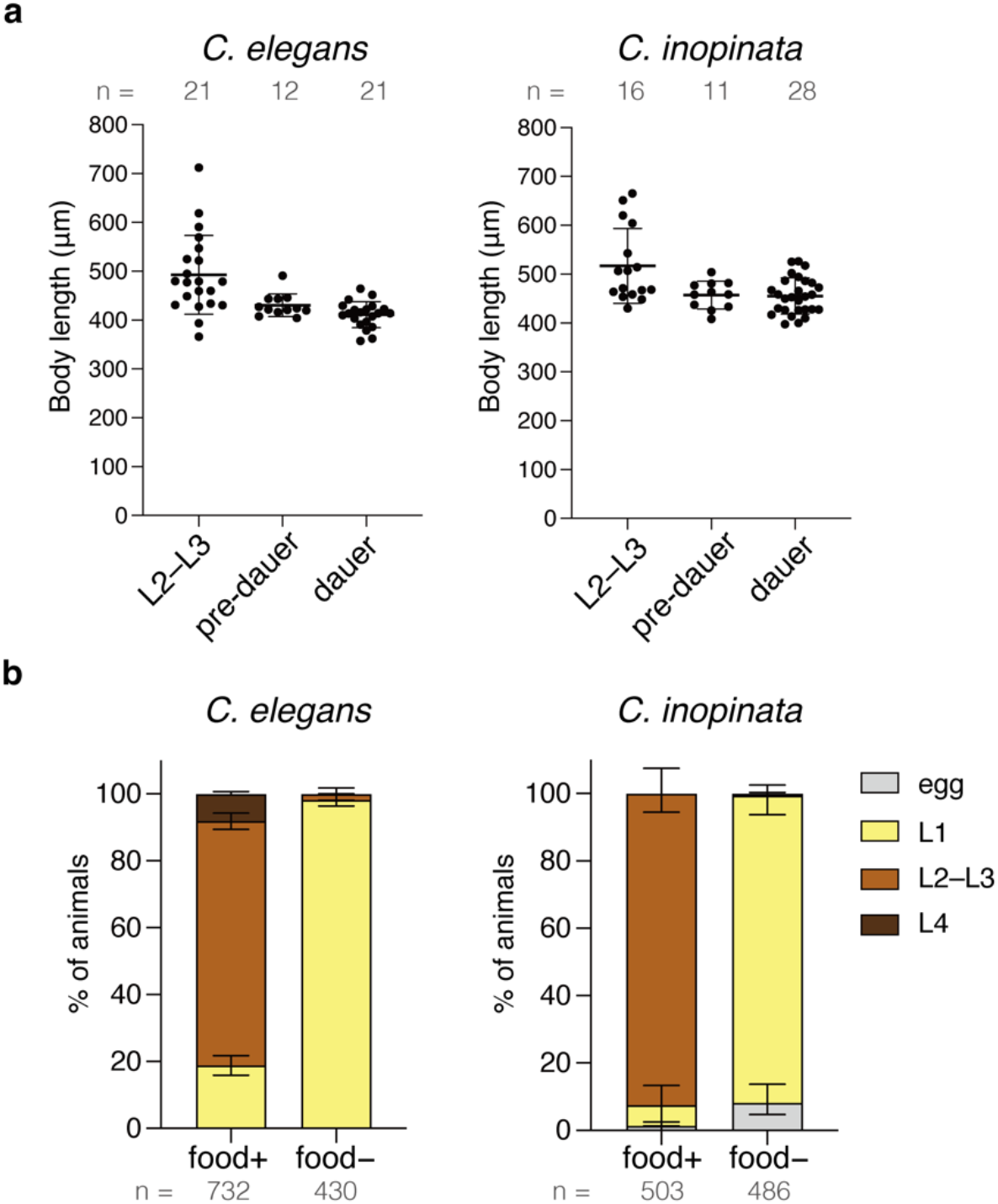
Preparation of pre-dauer and starved L1 larvae in *C. elegans* and *C. inopinata*. a) Body length of the L2–L3, pre-dauer, and dauer larvae in the *Cel/Cin-col-183p::mCherry* strains. The body length of pre-dauer larvae is similar to that of dauer larvae. Each dot represents an individual animal. b) Developmental stages of the *Cel/Cin-col-183p::mCherry* strains under fed (food+) and starvation (food−) conditions. Data were collected ∼36 h after plating gravid adults for *C. elegans* and ∼48 h for *C. inopinata*. In both strains, worms arrested at the L1 stage under starvation conditions. Stacked bar graphs show the proportion of animals at each developmental stage: egg (gray), L1(yellow), L2–L3 (brown), and L4 (dark brown). At least 50 animals were analyzed per plate, with 3–6 plates examined. In a) and b), n = the total number of animals analyzed. Bars indicate mean ± SD.

**Supplementary Fig. 2.**
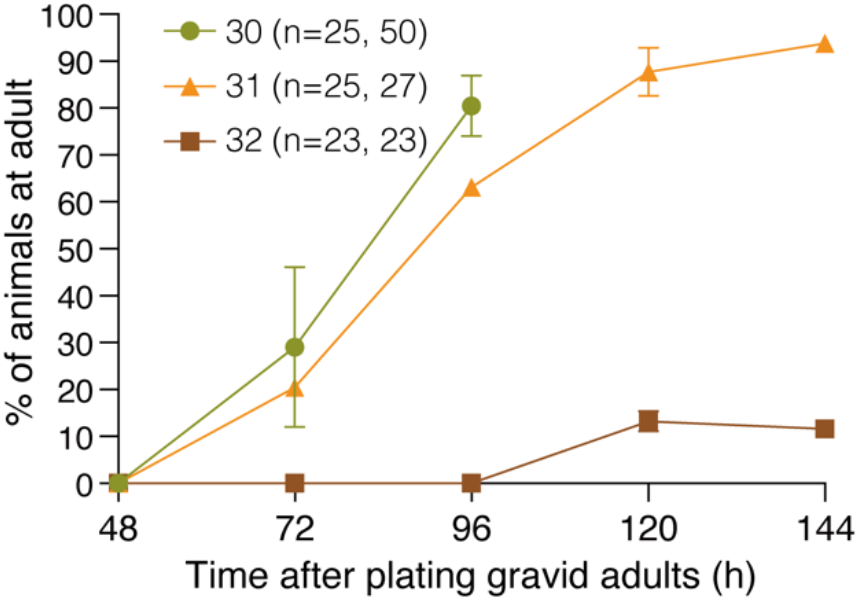
Determination of the high-temperature condition for *C. inopinata*. Proportion of F1 animals reaching adulthood over time after plating gravid P0 adults of the wild-type *C. inopinata* at 30°C (green), 31°C (orange), and 32°C (brown). Most animals at 30°C and 31°C developed to adulthood, whereas most animals at 32°C did not reach adulthood. Measurements at 30°C were terminated after 120 h because numerous next-generation larvae had appeared. Based on these results, 31°C was used as the high-temperature condition in this study. Each condition consisted of 2 plates, n = the number of animals analyzed. Bars indicate mean ± SD.

**Supplementary Fig. 3.**
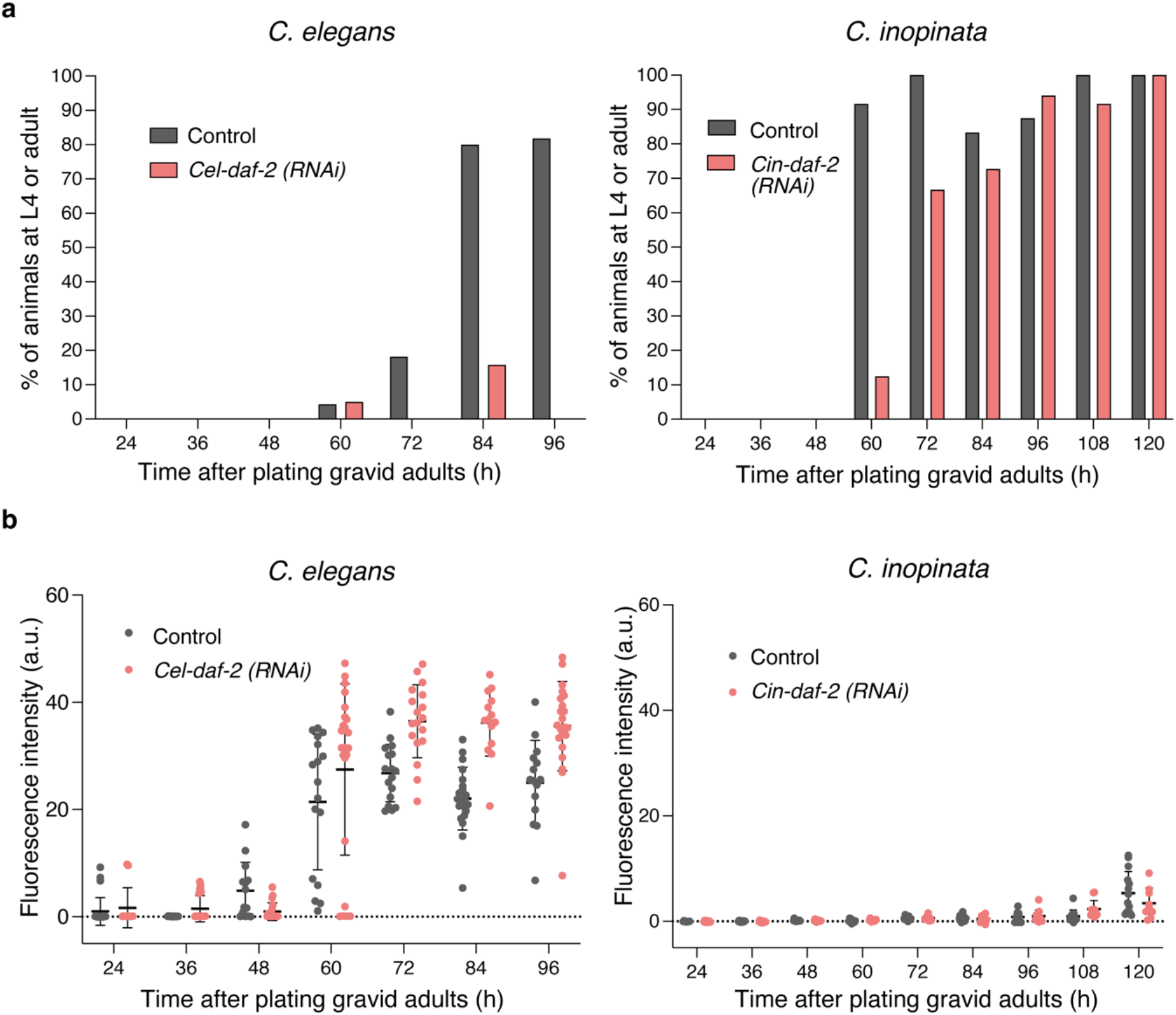
Time-course analysis of responses to *daf-2* RNAi in *C. elegans* and *C. inopinata*. a) Time course of proportion of animals reaching the L4 or adult stage, measured every 12 h at 27°C, which corresponds to a high temperature condition for *C. elegans* and an optimal temperature for *C. inopinata*, under control and *daf-2* RNAi conditions. b) Time course of mCherry fluorescence intensity (arbitrary units, a.u.) in individual animals from the same plates as in (a). Based on these results, 72 h at 27°C was selected for the analysis shown in Fig. 3. Each dot represents an individual animal. In a) and b), a single plate containing 8–25 animals was analyzed at each time point. Bars indicate mean ± SD.

**Supplementary Table 1.** Strain list.

| Species | Strain | Genotype | Culture temperature | Reference |
| --- | --- | --- | --- | --- |
| *C. elegans* | N2 | Wild type | 20°C | CGC |
| *C. elegans* | SA1692 | *tjTi122[Cel-col-183p::mCherry::Cel-unc-54 3'UTR, Cel-rps-0p::HygR::Cel-unc-54 3'UTR]I* | 23.5°C | This study |
| *C. elegans* | SA1696 | *tjTi126[Cel-ets-10p::gfp::Cel-unc-54 3'UTR, Cel-rps-0p::HygR:: Cel-unc-54 3'UTR]V* | 23.5°C | This study |
| *C. inopinata* | NKZ35 | Wild type | 27°C | Kanzaki *et al.*, 2018 |
| *C. inopinata* | SA1660 | *tjIs376[Cin-col-183p::mCherry::Cel-unc-54 3'UTR; Cel-rps-0p::HygR::Cel-unc-54 3'UTR], [Cin-ets-10p::gfp::Cel-unc-54 3'UTR; Cel-rps-0p::HygR::Cel-unc-54 3'UTR]* | 27°C | This study |
| *C. inopinata* | SA1664 | *tjIs377[Cin-col-183p::mCherry::Cel-unc-54 3'UTR; Cel-rps-0p::HygR:: Cel-unc-54 3'UTR]* | 27°C | This study |

**Supplementary Table 2.**
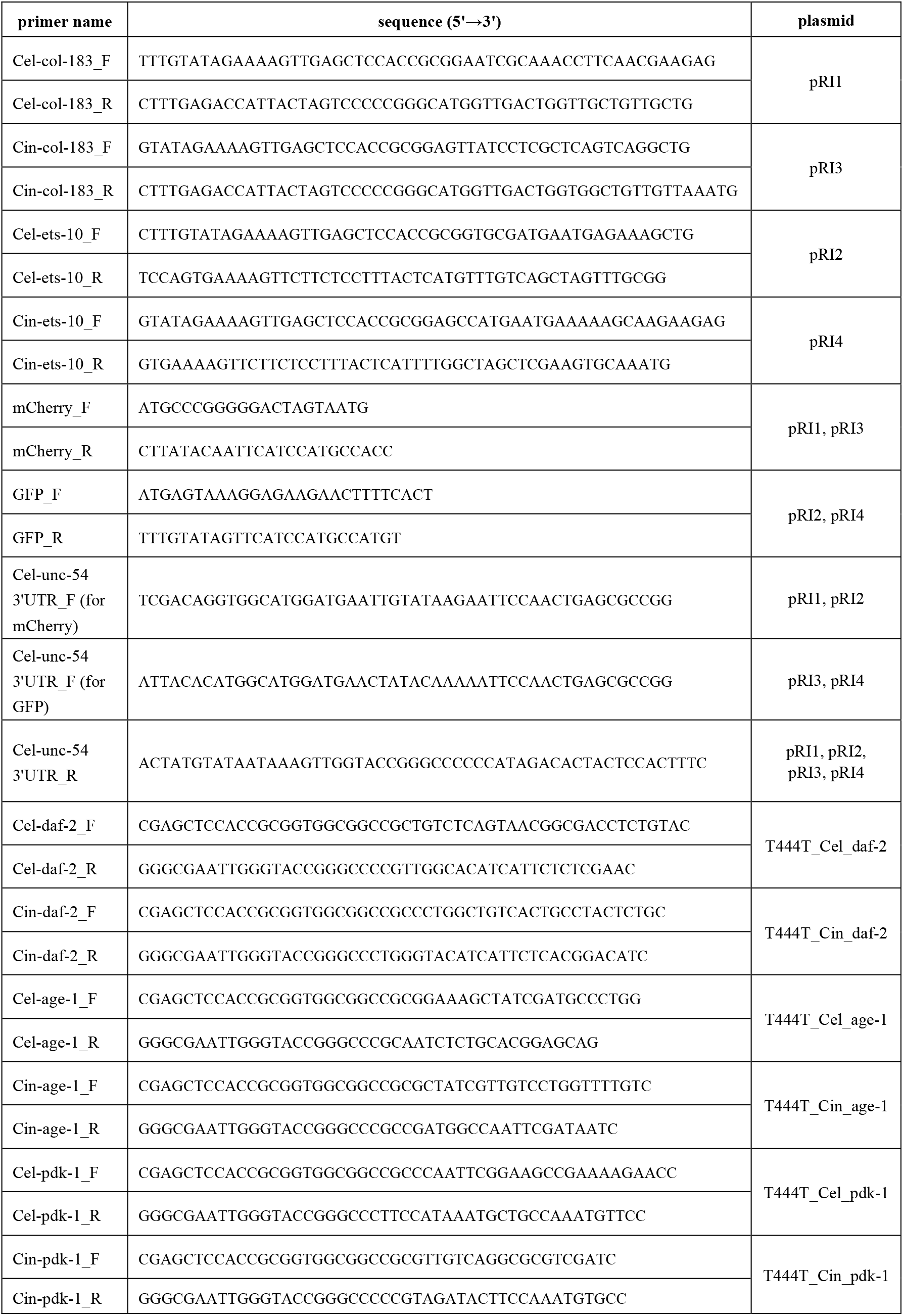
Primer list.

## References

Ailion, M., Thomas, J.H., 2000. Dauer Formation Induced by High Temperatures in Caenorhabditis elegans. Genetics 156, 1047–1067. 10.1093/genetics/156.3.1047

Brenner, S., 1974. The genetics of Caenorhabditis elegans. Genetics 77, 71–94. 10.1093/genetics/77.1.71

Byerly, L., Cassada, R.C., Russell, R.L., 1976. The life cycle of the nematode Caenorhabditis elegans: I. Wild-type growth and reproduction. Dev. Biol. 51, 23–33. 10.1016/0012-1606(76)90119-6

Cassada, R.C., Russell, R.L., 1975. The dauerlarva, a post-embryonic developmental variant of the nematode Caenorhabditis elegans. Dev. Biol. 46, 326–342. 10.1016/0012-1606(75)90109-8

Dillin, A., Crawford, D.K., Kenyon, C., 2002. Timing Requirements for Insulin/IGF-1 Signaling in C. elegans. Science 298, 830–834. 10.1126/science.1074240

Félix, M.-A., Braendle, C., 2010. The natural history of Caenorhabditis elegans. Curr. Biol. 20, R965–R969. 10.1016/j.cub.2010.09.050

Fielenbach, N., Antebi, A., 2008. C. elegans dauer formation and the molecular basis of plasticity. Genes Dev. 22, 2149–2165. 10.1101/gad.1701508

Frøkjær-Jensen, C., Davis, M.W., Sarov, M., Taylor, J., Flibotte, S., LaBella, M., Pozniakovsky, A., Moerman, D.G., Jorgensen, E.M., 2014. Random and targeted transgene insertion in Caenorhabditis elegans using a modified Mos1 transposon. Nat. Methods 11, 529–534. 10.1038/nmeth.2889

Gibson, D.G., Young, L., Chuang, R.-Y., Venter, J.C., Hutchison, C.A., Smith, H.O., 2009. Enzymatic assembly of DNA molecules up to several hundred kilobases. Nat. Methods 6, 343–345. 10.1038/nmeth.1318

Gilabert, A., Curran, D.M., Harvey, S.C., Wasmuth, J.D., 2016. Expanding the view on the evolution of the nematode dauer signalling pathways: refinement through gene gain and pathway co-option. BMC Genomics 17, 476. 10.1186/s12864-016-2770-7

Golden, J.W., Riddle, D.L., 1982. A Pheromone Influences Larval Development in the Nematode Caenorhabditis elegans. Science 218, 578–580. 10.1126/science.6896933

Golden, J.W., Riddle, D.L., 1984. The Caenorhabditis elegans dauer larva: Developmental effects of pheromone, food, and temperature. Dev. Biol. 102, 368–378. 10.1016/0012-1606(84)90201-X

Hammerschmith, E.W., Woodruff, G.C., Moser, K.A., Johnson, E., Phillips, P.C., 2022. Opposing directions of stage-specific body shape change in a close relative of C. elegans. BMC Zool. 7. 10.1186/s40850-022-00131-y

Johnson, T.E., Mitchell, D.H., Kline, S., Kemal, R., Foy, J., 1984. Arresting development arrests aging in the nematode Caenorhabditis elegans. Mech. Ageing Dev. 28, 23–40. 10.1016/0047-6374(84)90150-7

Kamath, R.S., Martinez-Campos, M., Zipperlen, P., Fraser, A.G., Ahringer, J., 2000. Effectiveness of specific RNA-mediated interference through ingested double-stranded RNA in Caenorhabditis elegans. Genome Biol. 2, RESEARCH0002. 10.1186/gb-2000-2-1-research0002

Kanzaki, N., Tsai, I.J., Tanaka, R., Hunt, V.L., Liu, D., Tsuyama, K., Maeda, Y., Namai, S., Kumagai, R., Tracey, A., Holroyd, N., Doyle, S.R., Woodruff, G.C., Murase, K., Kitazume, H., Chai, C., Akagi, A., Panda, O., Ke, H.-M., Schroeder, F.C., Wang, J., Berriman, M., Sternberg, P.W., Sugimoto, A., Kikuchi, T., 2018. Biology and genome of a newly discovered sibling species of Caenorhabditis elegans. Nat. Commun. 9, 3216. 10.1038/s41467-018-05712-5

McSorley, R., 2003. Adaptations of nematodes to environmental extremes. Fla. Entomol. 86, 138–142. 10.1653/0015-4040(2003)086%5B0138:AONTEE%5D2.0.CO;2

Mörck, C., Pilon, M., 2006. C. elegans feeding defective mutants have shorter body lengths and increased autophagy. BMC Dev. Biol. 6, 39. 10.1186/1471-213X-6-39

Nguyen, C.Q., Hall, D.H., Yang, Y., Fitch, D.H.A., 1999. Morphogenesis of the Caenorhabditis elegans Male Tail Tip. Dev. Biol. 207, 86–106. 10.1006/dbio.1998.9173

Oomura, S., Tsuyama, K., Haruta, N., Sugimoto, A., 2022. Transgenesis of the gonochoristic nematode Caenorhabditis inopinata by microparticle bombardment with hygromycin B selection. MicroPublication Biol. 10.17912. 10.17912/micropub.biology.000564

Paradis, S., Ailion, M., Toker, A., Thomas, J.H., Ruvkun, G., 1999. A PDK1 homolog is necessary and sufficient to transduce AGE-1 PI3 kinase signals that regulate diapause in Caenorhabditis elegans. Genes Dev. 13, 1438–1452. 10.1101/gad.13.11.1438

Roeber, F., Jex, A.R., Gasser, R.B., 2013. Advances in the diagnosis of key gastrointestinal nematode infections of livestock, with an emphasis on small ruminants. Biotechnol. Adv. 31, 1135–1152. 10.1016/j.biotechadv.2013.01.008

Shih, P.-Y., Lee, J.S., Sternberg, P.W., 2019. Genetic markers enable the verification and manipulation of the dauer entry decision. Dev. Biol. 454, 170–180. 10.1016/j.ydbio.2019.06.009

Sturm, Á., Saskői, É., Tibor, K., Weinhardt, N., Vellai, T., 2018. Highly efficient RNAi and Cas9-based auto-cloning systems for C. elegans research. Nucleic Acids Res. 46, e105– e105. 10.1093/nar/gky516

Timmons, L., Fire, A., 1998. Specific interference by ingested dsRNA. Nature 395, 854–854. 10.1038/27579

Toya, M., Iida, Y., Sugimoto, A., 2010. Imaging of mitotic spindle dynamics in Caenorhabditis elegans embryos. Methods Cell Biol. 97, 359–372. 10.1016/S0091-679X(10)97019-2

Vlaar, L.E., Bertran, A., Rahimi, M., Dong, L., Kammenga, J.E., Helder, J., Goverse, A., Bouwmeester, H.J., 2021. On the role of dauer in the adaptation of nematodes to a parasitic lifestyle. Parasit. Vectors 14, 554. 10.1186/s13071-021-04953-6

Woodruff, G.C., Johnson, E., Phillips, P.C., 2019. A large close relative of C. elegans is slow-developing but not long-lived. BMC Evol. Biol. 19, 74. 10.1186/s12862-019-1388-1

Zhao, L., Zhang, S., Wei, W., Hao, H., Zhang, B., Butcher, R.A., Sun, J., 2013. Chemical Signals Synchronize the Life Cycles of a Plant-Parasitic Nematode and Its Vector Beetle. Curr. Biol. 23, 2038–2043. 10.1016/j.cub.2013.08.041

